# New purifications reveal yeast and human stress granule cores are discrete particles with complex transcriptomes and proteomes

**DOI:** 10.1101/2023.10.12.562116

**Authors:** Natalia A. Demeshkina, Adrian R. Ferré-D’Amaré

## Abstract

Stress granules are a conserved response of eukaryotic cells to environmental insults. These cytoplasmic ribonucleoprotein condensates have hitherto been primarily studied by microscopy, which showed previously that they comprise dense ≈200 nm cores embedded in a diffuse shell. We have developed new purifications of budding yeast and mammalian (HEK293T cell) stress granule cores that do not rely on immunoprecipitation of candidate protein constituents. These unbiased preparations reveal that stress granule cores are discrete particles with variable size (mean, 135 nm and 225 nm, yeast and human, respectively) and shape. Proteomics and transcriptomics demonstrate complex composition. The results of hybrid chain reaction FISH analyses in HEK293T cells are consistent with stress granule cores having heterogeneous composition, *i.e*., each stress granule core particle contains only a limited number of mRNA species. Biochemical purification now opens the way to mechanistic analysis of the heterogeneity and complexity of stress granules.

## INTRODUCTION

When eukaryotic cells suffer environmental insults, such as oxidative and osmotic stress, nutrient deprivation, UV irradiation and heat shock, they universally respond by cessation of most protein synthesis. Then, their translationally inactive mRNAs associate with other cellular components into cytoplasmic assemblies called stress granules^1–4^. Upon stress alleviation, these ribonucleoprotein (RNP) granules dissipate, and the released mRNAs re-engage in translation^5, 6^. Stress granule formation and disassembly, which are conserved throughout eukaryotes, are thought to be cytoprotective^7^. Many pathways converge into turnover of stress granules^7–11^, whose dysregulation is linked to a variety of disease states, including neurodegeneration and cancer^12, 13^. Stress granules are also critical in anti-viral response^14^, and some viruses hijack them as part of their proliferation strategy^15^.

Stress granules were first discovered by electron microscopy of heat-shocked tomato cells^16^. Subsequent studies demonstrated their formation in response to other stressors in diverse eukaryotic cells. In most studies, stress granules have been imaged by light microscopy, relying on fluorescently labeled RNA and protein factors identified as components of the granules^17–19^. Together, electron and light microscopy studies indicate that stress granules are comprised of a denser core with an estimated diameter of 150-270 nm, surrounded by a more diffuse shell^6, 17, 20^.

The large size, lack of an enclosing bilayer membrane, and dynamic nature of stress granules have made their purification challenging^5, 6^. Pioneering studies employed immunoprecipitation of tagged proteins as the main enrichment step^17, 21, 22^. That approach benefitted from expediency^23^, accessible scale, and diversity of potential bait proteins, and provided information on RNA and protein components that are overrepresented in the fractionated material. The published protocols have, however, four substantive limitations. First, the relatively small scale of the reported purifications (which start with ≈ 50 mL yeast cultures or mammalian cells collected from a 500 cm^2^ growth^21^) produced correspondingly small quantities of enriched fractions, ultimately limiting the depth and redundancy of the transcriptomic and proteomic analyses. Second, the protocols entail cell lysis with a large excess of buffer, in the absence of stabilizing agents. These manipulations may lead to disruption of the structure of stress granules and stripping of integral components. Third, the protocols all recover material of interest from pellets, which are enriched in high mass, actively translating RNPs (*e.g.,* polysomes). Fourth and most critically, because the bait proteins exist both, in the cytosol and in stress granules, the transcriptomic and proteomic results are confounded by components that associate with the bait (*e.g*., poly(A) binding protein (PABP), G3BP1/2) in the cytosolic compartment. Immunoprecipitation of cellular lysates without robust orthogonal fractionation will result in enrichment of any particles, regardless size, that are decorated by the bait.

We have now developed new purifications of stress granule cores from budding yeast and human embryonic kidney 293T (HEK293T) cells that are unbiased by assumptions on the subcellular distribution of specific candidate proteins. Our protocols do not rely on immunoprecipitation. Instead, we exploit the density and hydrodynamic radius of stress granule cores, which are independent of the localization or immunoreactivity of bait proteins. We adapted lysis conditions previously employed successfully for the isolation of large, delicate and dynamic RNP complexes (*e.g.*, spliceosomes), and never pelleted fractions of interest, to exclude polysomes. Our purifications start from much larger cell culture volumes (≈ 100 x) than those previously reported, providing ample material for high-redundancy proteomics and transcriptomics. The extensive datasets of proteins and RNAs resulting from our purified yeast and mammalian stress granule cores will constitute a valuable resource for further genetic, biochemical and cell-biological analyses of these large cytoplasmic assemblies that are a key adaptation mechanism in all eukaryotic cells.

## RESULTS

### Purification of yeast and mammalian stress granule cores

We developed a purification of *Saccharomyces cerevisiae* stress granule cores formed in response to sodium azide treatment^24^ (Figure 1), which causes oxidative stress by inhibiting respiration. Unlike the published procedure^17, 21^, which employed centrifugal pelleting followed by immunoprecipitation (Figure S1A), our approach was designed to isolate stress granule cores based on their average size (∼ 180 nm), as determined previously^17^ by *in vivo* imaging in yeast. To preserve the integrity of large assemblages, cell lysis was performed without detergents, under osmotic conditions imitating the dense cytosol milieu (Figure 1A). Stress granule cores were always collected from soluble fractions and never from pellets, by following green fluorescent protein (GFP) fusions of the stress granule markers^17^ eIF4G1 or PABP, which bind to the 5’ and 3’-untranslated regions of mRNAs, respectively. To remove translationally active mRNAs (*i.e*., in polysomes) from other cytoplasmic complexes, we employed sucrose density-gradient centrifugation (SDGC) (Figure 1A). This revealed comparatively low-density, GFP-enriched fractions with large nucleic acids distinct from translationally active fractions as well as ribosomal subunits (Figures 1C and 1D). For comparison, we observed similar low-density, marker protein-GFP-fusion enriched fractions when we subjected to SDGC stress granule-enriched material obtained through the immunoprecipitation protocol of Jain et al.^17^ (Figures S1B-S1D). These fractions with large nucleic acids were markedly diminished in purifications treated prior to stress with cycloheximide (Figure S1D), which is widely used in stress granules studies because it inhibits their assembly^5, 17^.

**Figure 1.**
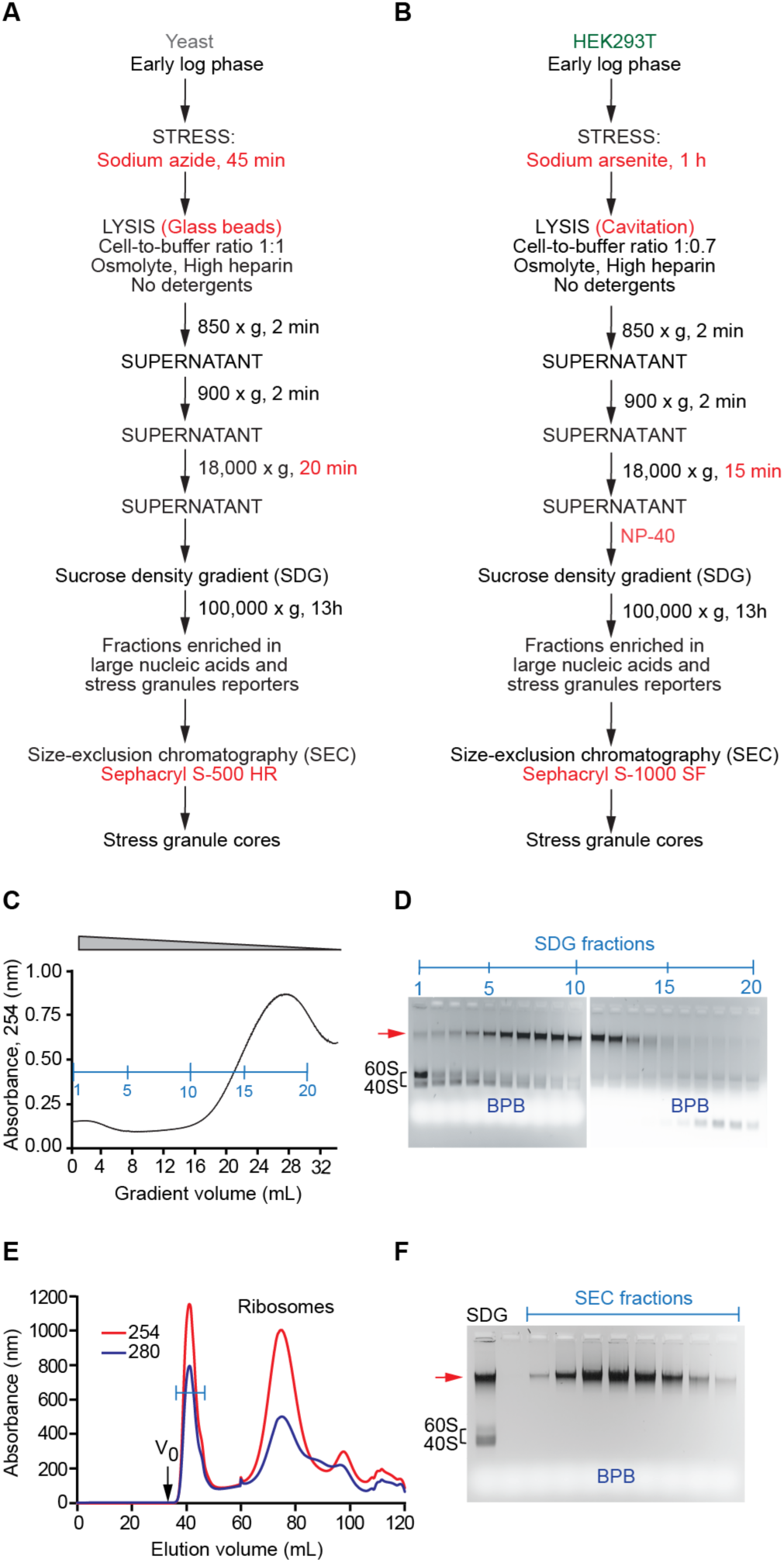
Purification of stress granule cores from yeast and mammalian cells. (**A**) Schematic of the sucrose density-gradient centrifugation, size-exclusion chromatography (SDGC-SEC) purification of stress granule cores from *S. cerevisiae* cells. (**B**) Schematic of the SDGC-SEC stress granule core purification from HEK293T cells. Differences between (A) and (B) indicated in red. (**C**) Sucrose density-gradient centrifugation profile of the cytoplasmic fraction of *S.* c*erevisiae* exposed to oxidative stress. Blue line, fractions. (**D**) Agarose gel electrophoretic analysis of fractions from (C). Arrow, electrophoretic mobility of high molecular weight species enriched in nucleic acids. Migration position of 40S and 60S ribosomal subunits and bromophenol blue (BPB) indicated. (**E**) SEC of combined fractions 2 – 12 from (C). Blue line, fractions. *V*_0_, void volume. (**F**) Agarose gel electrophoretic analysis of fractions from (E).

To achieve further purification on the basis of hydrodynamic radius, we next subjected the SDGC-fractionated material to size-exclusion chromatography (SEC), using a resin with a nominal exclusion limit of 100 MDa (Figure S2A). The chromatography was performed in the presence of dilute urea, to separate high molecular weight components from loosely bound larger and smaller cytoplasmic constituents (Figures 1E and 1F). The elution volume of the well-defined absorbance peak was consistently larger than the void volume (*V*_0_) of the column employed, implying that the material entered the resin, and indicative of discrete particles, rather than amorphous aggregates. To exclude peculiarities of a specific yeast genotype, we carried out purifications from three *S. cerevisiae* strains (YER165W with PABP-GFP, JD1370 and YAG1021), grown in either rich (YPDA) or in synthetic defined (SD) media (Methods). All purifications yielded biophysically indistinguishable SDGC-SEC fractions. Overall, our procedure allowed us to isolate large, translationally inert cytoplasmic particles, free from polysomes, which was not possible using the immunoprecipitation-based protocol (Figure S1E).

We next adapted our purification strategy to mammalian cells (Figure 1B). Our preparations each started with 8 L of HEK293T cells grown in suspension. Sodium arsenite is commonly used to subject mammalian cells to oxidative stress and induce stress granule formation^25^. We treated our HEK293T cells with sodium arsenite in early log phase (equivalent to ≈ 20% confluency), to avoid confounding effects of growth to very late log phase (≈ 80-90% confluency) as has been typical^17, 21^ for the published immunoprecipitation-based preparations. For lysis, we employed controlled cavitation (Methods). This method has been previously demonstrated to rupture the plasma membrane while maintaining integrity of the nuclear envelope and other cytoplasmic membranous structures^26^. We performed lysis with minimal dilution (Figure 1B, cell-to-buffer ratio 1:0.7), hence, minimizing destabilization of large, non-covalent cytoplasmic structures. Similarly to yeast purification protocol, cells were lysed in the presence of a stabilizing agent that further helped to prevent rupture of nuclei and stabilized cytoplasmic structures. Our mammalian purification protocol is robust, allowing reproducible recovery of large cytoplasmic particles with substantively comparable properties to the purified yeast stress granule cores from the SDGC and SEC steps (Figures S2B and S2C). Both yeast and mammalian stress granule cores were maximally enriched at gradient fractions containing 17-20% sucrose (w/v). Like the yeast particles, mammalian stress granule cores also eluted well after the void volume of the SEC resin, consistent with their discrete, particulate nature.

### Physical and chemical properties of stress granule cores

When examined by nanoparticle tracking analysis, purified yeast stress granule cores show a relatively narrow size distribution peaking at 135 nm (Figure 2A). This is consistent with the conclusions of previous structured illumination microscopy analyses^17^, which indicated their diameter, *in vivo*, to be 179.5 ± 28.9 nm. Similarly, nanoparticle tracking analysis of HEK293T cell stress granule cores showed that they have a relatively narrow distribution peaking at 225 nm (Figure 2B). Together, these solution analyses show that, consistent with their behavior during purification by size-exclusion chromatography (Figures 1E and S2C), stress granule cores are discrete but heterogeneous particles, not amorphous aggregates, and that mammalian stress granule cores are larger, on average, than their yeast counterparts.

**Figure 2.**
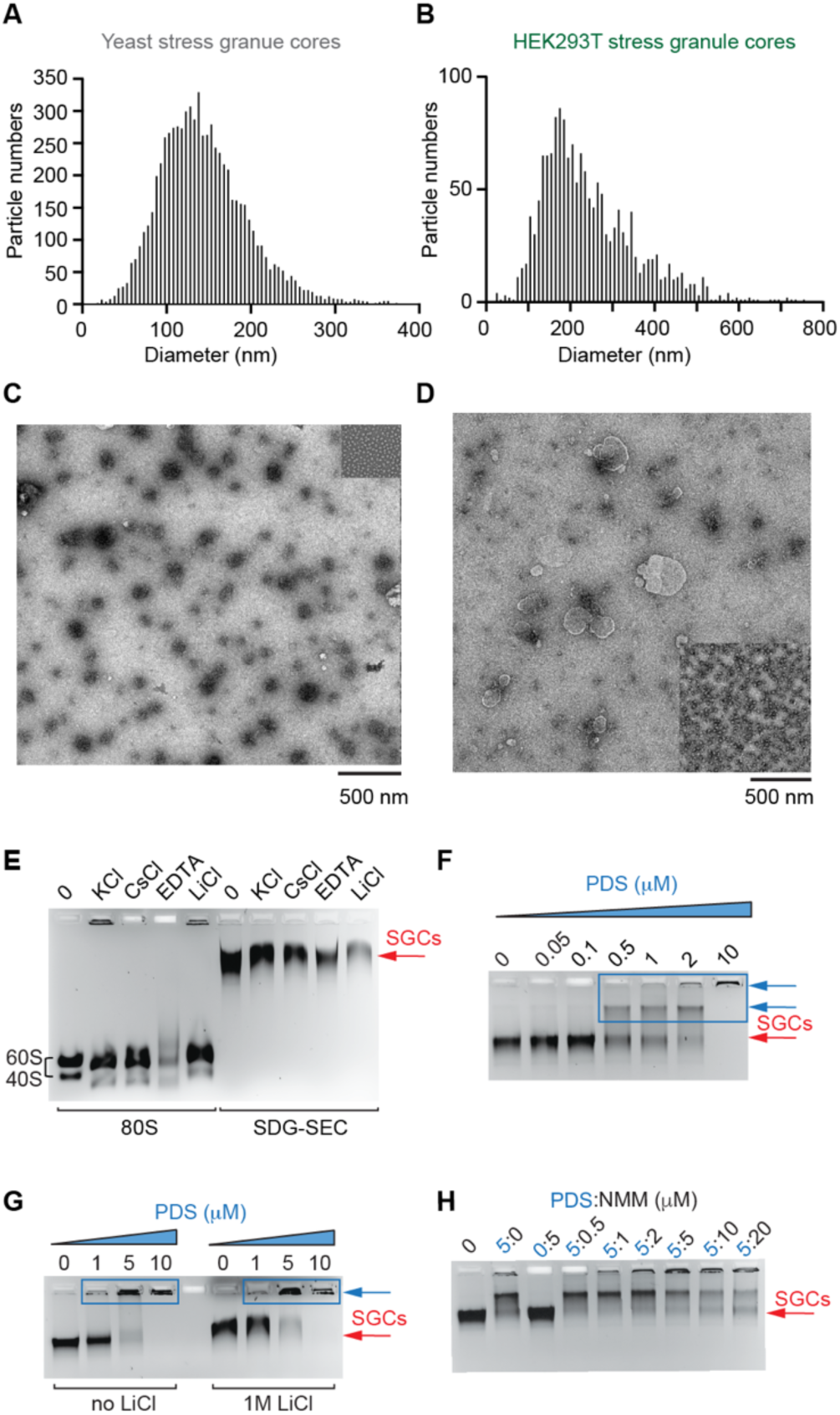
Physical and chemical characterization of SDGC-SEC stress granule cores. (**A**) Nanoparticle tracking analysis of yeast stress granule cores. The median particle size is 135 nm. (**B**) Nanoparticle tracking analysis of HEK293T stress granule cores. The median particle size is 225 nm. (**C**) Negative-stain electron micrograph of yeast stress granule cores; magnification (× 2,900). Inset, yeast 80S ribosomes at the same magnification. (**D**) Negative stain electron micrograph of HEK293T stress granule cores; magnification (× 2,900). Inset, pellet from sucrose-density gradient centrifugation step (Figure 1B) containing HEK293T polysomes and 80S ribosomes; same magnification. (**E**) Agarose gel electrophoretic analysis showing chemical stability of yeast stress granule cores and 80S ribosomes in the presence of 1M KCl, 1M CsCl, 50 mM EDTA or 1M LiCl. SEC, stress granule cores, SDC-SEC, sucrose density gradient-size exclusion-chromatography. (**F**) Interaction of yeast stress granule cores with pyridostatin (PDS), a G-quadruplex ligand. Blue arrow denotes stress granule cores fusion. (**G**) Effect of LiCl on PDS interaction with yeast stress granule cores. Blue arrow denotes stress granule cores fusion. (**H**) Interaction of yeast stress granule cores with the G-quadruplex ligands NMM and PDS.

Negative-stain transmission electron microscopy of purified yeast stress granule cores shows particles with a size distribution consistent with that detected in solution by nanoparticle tracking analysis, and with irregular but compact shapes (Figure 2C). Some of the observed shape variability could be the result of embedding the specimens in uranyl formate^27^. Comparison with purified yeast 80S ribosomes shows that stress granule cores are at typically ≈ 10-15 x larger in linear dimensions. During our purifications, we found that mammalian stress granule cores appeared to co-purify more than their yeast counterparts with cellular membranes. After treatment with dodecyl-β-maltoside (Methods), negatively stained HEK293T cell stress granule cores show a wider range in size distribution, compared to their yeast counterparts (Figure 2D). Overall, electron microscopy confirms that stress granule cores are large, discrete particles with variable shape and size.

Previous studies suggested^28, 29^ the involvement of G-quadruplexes in stress granule formation. We found that, similarly to what was reported earlier for immunoprecipitated material^17^, purified stress granule cores are relatively stable to incubation in high concentrations of the physiologic cation K^+^. However, the particles are strongly destabilized by LiCl (Figure 2E). Destabilization by Li^+^ is a hallmark of G-quadruplexes. We therefore tested the effects of two G-quadruplex-specific compounds, pyridostatin (PDS)^30^ and N-methyl mesoporphyrin IX (NMM)^31^. PDS induced dose-dependent fusion of purified stress granule cores (Figure 2F) that was not largely influenced by LiCl (Figure 2G). The PDS-induced aggregation was partially suppressed by NMM (Figure 2H). However, when the stress granule cores were preincubated with NMM, and PDS was later titrated, aggregation ensued (data not shown). The susceptibility of stress granule cores to Li^+^ and PDS is consistent with G-quadruplexes playing a structural role in them.

### Proteomes of purified yeast and mammalian stress granule cores

We subjected our purified yeast and mammalian stress granule cores to proteomic analysis (Methods). To compare the protein composition of the yeast stress granule cores from the new purifications to the previously reported immunoprecipitation-based protocol^17, 21^ we performed, in parallel, the SDGC-SEC purification and the immunoprecipitation-based protocol starting with the same yeast culture and cell mass (Figure 3A and Table S1), with identical downstream treatments. This experimental design avoids the pitfalls of comparing protein composition when strains and methods differ. This comparative analysis demonstrated a substantial overlap (45%) between the proteomes of the SGDC-SEC-purified particles and the immunoprecipitation-enriched material.

**Figure 3.**
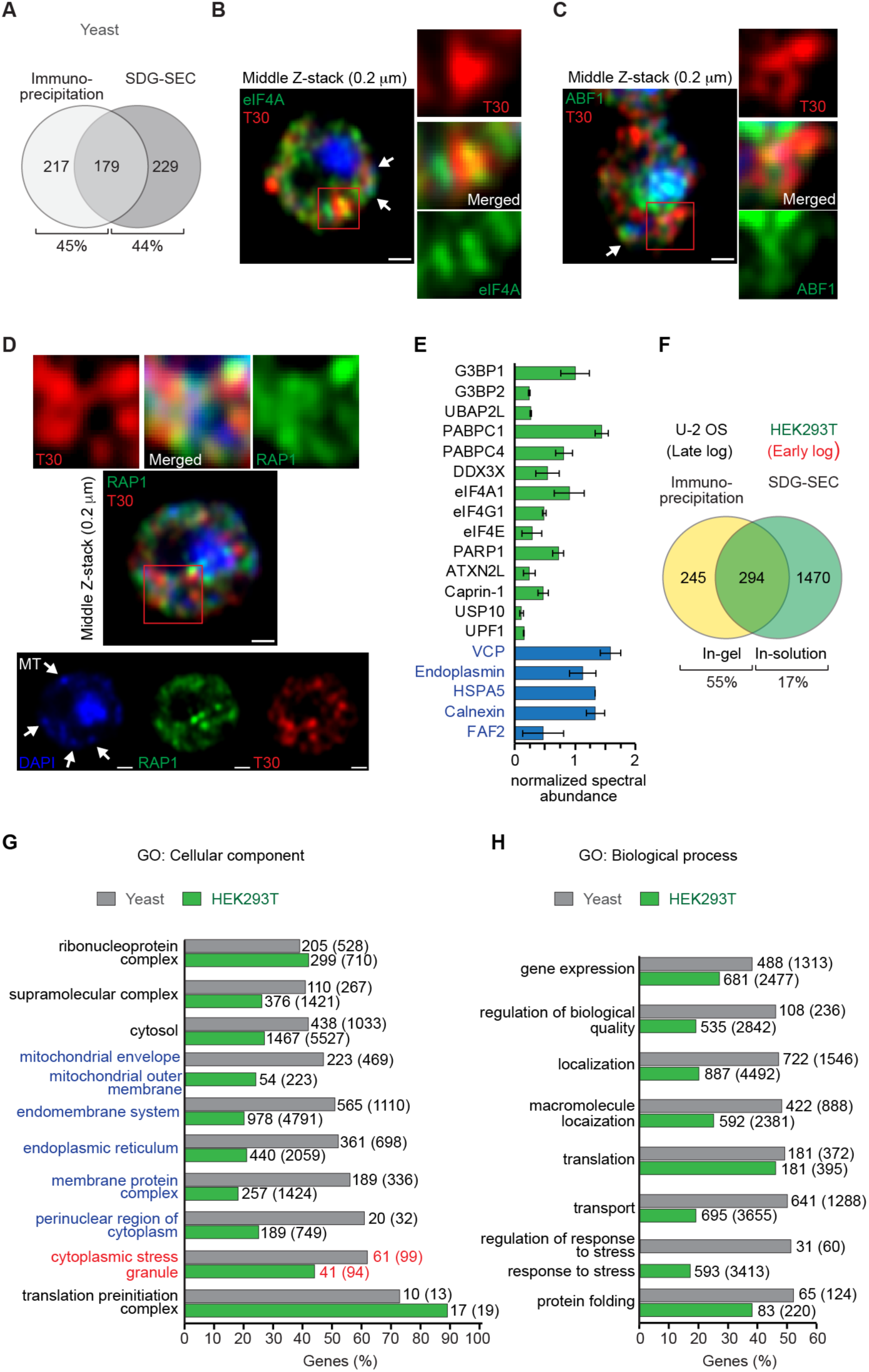
Proteomes of stress granule cores from yeast and mammalian cells. (**A**) Overlap between proteomes of yeast stress granules enriched by immunoprecipitation and yeast stress granule cores isolated by sucrose-density gradient-size-exclusion. The comparison was conducted in parallel (same cell amounts, identical downstream treatments) for a strain with an endogenous PAB1-GFP fusion. (**B**-**D**) Fluorescent microscopy of yeast colocalizes eIF4A (B), ABF1 (C) and RAP1 (D) with polyadenylated mRNAs (red) within stress granules. Green, GFP fusion. Blue, nucleus and mitochondrion (MT)^36^ stained by DAPI. Scale bar 0.5 μm. (**E**) Example proteins from purified HEK293Tstress granule cores (n=2; average ± SD). Relative abundance of proteins between datasets was normalized using G3BP1 (Methods). Black, previously characterized stress granule proteins. Blue, membrane-associated proteins. (**F**) Overlap between proteomes of U-2 OS mammalian stress granules enriched by immunoprecipitation^17^ and stress granule cores isolated from HEK293T cells (this work). (**G**, **H**) GO terms for proteins from stress granule cores of *S. cerevisiae* grown in minimal medium (n=2) and HEK293T cells (n=3). Selection criteria: (1) presence in both datasets; (2) abundant GO term; (3) high percentage within a GO term. Average number of genes and total genes (in parentheses) in each GO category are indicated. In (G), blue marks terms involved in membrane metabolism.

The proteomes of our purified particles include numerous proteins recognized as stress granule constituents, such as PABP, the eIF4F complex (comprised of eIF4A, eIF4G and eIF4E/CDC33), translational helicases DED1 and DHH1, eIF3s, heat-shock family SSA ATPases, chaperonin containing tailless (CCT) complex, tubulins, TMA19 and TYS1 (Table 1). In addition to invariant constituents, our analysis shows that other protein components of stress granule cores can vary. Thus, purified cores from cells grown in rich media lacked eIF2α (Tables 1 and S2); however, those purified from *S. cerevisiae* fermented in SD medium contained this factor (Tables 1 and S2). It should be noted that while phosphorylation of eIF2α triggers assembly of stress granules in mammalian cells^1^, this protein is not required for stress-granule formation in yeast^19^. Presence of PBP1, the yeast ortholog of mammalian Ataxin-2, was also depended on growth conditions in strain JD1370 (Tables 1 and S2), being present when cells were grown in minimal medium, but absent in rich media growths. In contrast, stress granule cores purified from the strain YER165, which carries an endogenous PABP-GFP fusion, grown in rich media exhibited abundant PBP1 (Table S2). These examples illustrate how some proteins considered to be markers can be absent from stress granule cores, probably reflecting, in addition to organism, strain, growth conditions, type and timing of stressor^19^ also the purification methods and downstream manipulations (*e.g*., in-gel versus in-solution protein digestion for MS analysis), emphasizing the importance of controlled, parallel purifications for comparative analyses.

Our proteomic analyses show that numerous proteins not directly linked to the stress response or translation are present in yeast stress granule cores. Thus, for instance, the RNA-helicase DBP5, which is a constituent of RNA export granules^32^, was consistently present in our purifications (Table S2). This essential and conserved RNA helicase is primarily active on the cytoplasmic side of the nuclear envelope, which is continuous with the endoplasmic reticulum^33^. The latter often contacts stress granules. All our preparations also contained proteins involved in transcription and DNA metabolism, such as ABFs and RAP1. Components of the minichromosomal maintenance (MCM) complex and RVB1,2 that were previously identified in the proteomes of stress granules enriched by immunoprecipitation^17^ were also present in our purified cores (Tables 1 and S2), suggesting that they are not contaminants, but authentic constituents. Overall, our analysis considerably extends the list of yeast stress granule core proteins (Table S2).

We analyzed the localization of ABF1 and RAP1, two of the newly identified yeast stress granule core components, by fluorescent microscopy. Yeast strains with the chromosomally encoded fusions ABF1-GFP and RAP1-GFP, as well as a T30 probe for the poly-A tails of mRNAs (Figures 3B-3D) were employed. This showed that ABF1 (Figure 3C) and RAP1 (Figure 3D) colocalize with the signal for mRNA in stress granules in a manner similar to eIF4A-GFP (Figure 3B). The latter is a key conserved stress granule regulator^34^, which is a part of the eIF4F complex. The presence in stress granules of the small GTPase RAP1 is consistent with its reported^35^ re-localization to the cytoplasm upon hypoxia. One notable aspect of the proteome of our purified stress granule cores is the presence of proteins associated with mitochondria. Staining of yeast cells with DAPI allows visualization of both, nuclei and mitochondria^36^ (Figure 3D). In many of our images, stress granules as demarcated by T30, eIF4A, ABF1 and RAP1 signals colocalized with mitochondria (Figure 3G), indicating that stress granules can closely associate with the respiratory organelle.

Proteomic analysis of purified mammalian stress granule cores confirmed the presence of numerous expected proteins, including G3PBs, the eIF4F complex, PABPCs, DDX3X, Ataxin-2-like protein and UBAP2L (Figure 3E and Table 1). Comparison with the previously reported proteome of immunoprecipitated stress granules from U-2 OS human osteosarcoma-derived cells stressed at late log phase, shows more than 50% overlap, even though different cell lines and purification procedures are involved (Figure 3F and Table S2 in Jain et al. 2016). Although at a modest titer, our HEK293T stress granule cores contain (Table S3) some components of the TREX complex (*e.g*., THOC2, THOC3, ALYREF and DDX39B), which function^37, 38^ at the interface of transcription, mRNA processing and mRNP nuclear export. No enrichment was present of major components of processing bodies (P-bodies), which are mainly involved in mRNA degradation^39, 40^. Similarly, the purified yeast stress granules also lacked P-body proteins (Table S2). An exception was the DEAD-box helicase DDX6 (DHH1 in yeast) (Table 1), which is essential in microRNA-induced gene silencing^41^, and has been reported to localize to both, P-bodies and stress granules. Overall, the proteomes of yeast and mammalian stress granule cores have many commonalities between them, indicative of phylogenetic conservation of this cytoplasmic response to environmental insults.

Gene ontology analyses of the proteomes of purified yeast and mammalian stress granule cores highlight many proteins in categories associated with cytoplasmic membrane systems including the mitochondrial network (Figures 3G and 3H). Earlier electron microscopic studies^20^ had noted a tight association of mammalian stress granules with mitochondria, and stress granules are known^42^ to regulate fatty acid metabolism by modulating activity of a voltage-dependent anion channel (VDAC). This channel, which is present in both of our stress granule core datasets (Figure S3 and Tables S2 and S3), mainly localizes to the outer mitochondrial membrane^43, 44^, and is a critical component of the dynamic interface between mitochondria and the endoplasmic reticulum, which governs homeostasis and cell death^45, 46^. Other examples of membrane-associated proteins are the AAA+ valosin containing protein (VCP or p97 in mammals, CDC48 in yeast^8, 47, 48^), FAF2 (ref. ^48^) and HSPA5 (ref. ^49^) (Figure 3E). The mammalian stress granule cores are also enriched in endoplasmin (HSP90B1, GP96 or GRP94), a key player in innate and adaptive immunity^50^, and Calnexin (IP90), a endoplasmic reticulum chaperone^51^ (Figure 3E). Possibly because of their role in organellar or granule traffic^17^, the purified stress granule cores also contain components of the cytoskeletal apparatus (Table 1 and Figure S3). Overall, association analysis of the proteomes of stress granule cores (Figures S3B and S3C) reveals them enmeshed in a complex network pathways, consistent with their being^7^ regulatory hubs of cellular homeostasis.

### RNA composition of yeast and mammalian stress granule cores

We next characterized the RNA composition of purified yeast stress granule cores. As expected, the majority of identified transcripts were mRNAs (Figure 4A), even though our analysis also showed the presence of numerous, comparatively rare, long non-coding RNAs (lncRNAs; Figure 4A). As our sequencing methodology did not explicitly search for short RNAs, their abundance may be underrepresented in our dataset. Previous studies^22^ of immunoprecipitated stress granules also reported some lncRNAs. We find that the majority of lncRNAs originate from the antisense strands of protein coding genes (Figure 4B).

**Figure 4.**
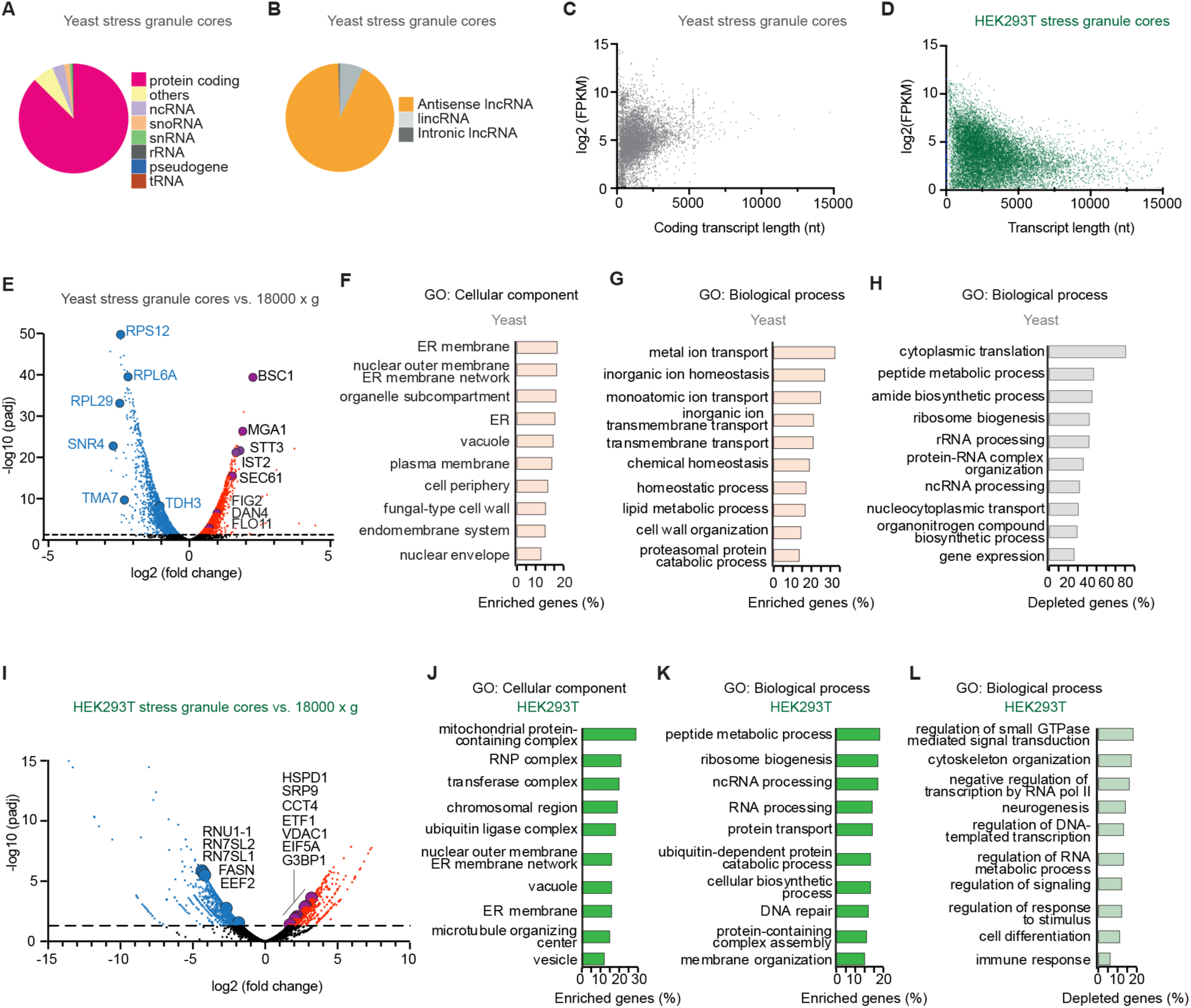
Transcriptomes of yeast and mammalian stress granule cores. (**A**) Distribution of identified RNA types within yeast stress granule cores (JD1370, rich medium). ‘Others’ category includes lncRNAs with *in silico* coding potential. (**B**) Major types of lncRNAs from yeast stress granule cores (JD1370, rich medium). (**C**) Length distribution of coding RNA transcripts from yeast stress granule cores (JD1370, rich medium). (**D**) Length distribution of RNA transcripts from HEK293T stress granule cores. (**E**) Volcano plot differential analysis for yeast stress granule cores and 18000 × g fraction (Figure 1A). Strain JD1370, minimal medium. (**F**-**H**) GO terms for enriched and depleted genes (n=2; p-adjusted < 0.05) for yeast stress granule core transcriptome. Selection criteria: abundant GO term (> 100 genes) and highest percentage of differentially expressed genes within a GO term. (I) Volcano plot differential analysis for HEK293T stress granule cores and 18000 × g fraction (Figure 1B). (**J**-**L**) GO terms for enriched and depleted genes (n=2; p-adjusted < 0.05) for HEK293T stress granule core transcriptome. Selection criteria: abundant GO term (> 300 genes) and highest percentage of differentially expressed genes within a GO term. In (E, I) example enriched (red) and depleted (blue) genes are highlighted.

The mRNAs isolated from our purified yeast stress granule cores had a broad length distribution, with an average of ∼1.4 kb (Figure 4C and Table S4), which is comparable to the average length^52, 53^ of yeast mRNAs (1.25-1.5 kb). Analysis of the RNA composition of purified HEK293T cell stress granule cores showed a wide length distribution, with an average of ∼2.3 kb (Figure 4D and Table S6). This is close to the average length of human coding transcripts. Our results contrast with those from previous analyses of mRNAs from pelleted and immunoprecipitated stress granule preparations from yeast and mammalian cells that found average RNA lengths of ∼2.7 and ∼7 kb, respectively. In that study, it was proposed that longer mRNAs preferentially form stress granules^22^. More recently, however, analysis of yeast stress granules where mRNA was characterized from all fractions of lysates (supernatants and pellets) obtained by sequential centrifugation without immunoprecipitation suggested that the propensity of yeast mRNAs to enter stress granules is not length-dependent^54^. The mRNA lengths we find are consistent with the latter study, and may indicate that use of bait proteins in immunoprecipitation approaches biases the enriched fractions to particles containing longer mRNAs. Alternatively, the mRNA composition of stress granules may differ between cells stressed in early *vs.* late log phase, or other experimental parameters.

To determine what mRNAs from the cellular pool are enriched in yeast stress granule cores, we compared mRNAs from the purified particles with those from the supernatant fraction prior to sucrose density-gradient centrifugation (Figure 1A). Classification of ∼440 enriched mRNAs according to gene ontology categories (Figures 4E-4H and Table S5) shows enhancement of genes associated with intracellular membranes, in particular the endoplasmic reticulum (Figure 4F). For instance, the products of genes SEC61, STT3 and IST2 (Figure 4E), play essential roles in the translocon channel, the oligosaccharyltransferase complex, and the endoplasmic reticulum-plasma membrane network, respectively. Another enriched set includes BSC1, whose product is part of a network of adaptation to oxidative stress^55^, and MGA1, a heat shock transcription factor whose loss results in decreased anaerobic growth^35^. We find that two most enriched mRNAs identified by immunoprecipitation^22^, FIG2 and DAN4, are also enriched in our datasets (Figure 4E). Gene ontology classes overrepresented in transcripts depleted from yeast stress granule cores are related to ribosomes and translation, rRNA processing and ribosomal biogenesis, and extend to amide biosynthetic processes (Figure 4H).

We performed similar analysis of mRNA enrichment in mammalian stress granule cores by comparing the transcriptomes of the purified particles and of the fraction prior to the sucrose density-gradient centrifugation step (Figure 1B). Some 2200 transcripts were significantly (p < 0.05) enriched in stress granule cores (Figure 4I and Table S7). Gene ontology analysis related these transcripts to multiple categories ranging from RNP and transferase complexes (Figures 4J) to various RNA processing pathways, and to DNA repair (Figures 4K). As in the case of yeast (Figure 4F), many were in classes involved in membrane networking (Figure 4J). Salient depleted categories included regulation of transcription, cell differentiation, immune response and neurodegeneration (Figure 4L). Overall, however, the enriched and depleted mRNAs did not sort into a clear set of categories, indicating that, as in yeast, mammalian stress granule cores comprise mRNAs from diverse pathways.

### Stress granule cores have heterogeneous mRNA composition

In addition to providing access to the proteomes and transcriptomes of yeast and mammalian stress granule cores, our purification reveals that these cytoplasmic constituents, hitherto characterized predominantly by fluorescent imaging methods, are discrete particles with a relatively low density, narrow size distribution and variable shape (Figures 2A-2D). The mean size and density of stress granule cores places physical constraints on their composition. Making the simplifying assumption of a spherical shape, the mean volume of one yeast stress granule core with a 50 nm radius is 5.2 x 10^-16^ cm^3^. The sucrose concentration at the peak of their distribution is ∼19% (w/v), with a density of 1.081 g•cm^3^, which yields a mean mass of 5.6 x 10^-16^ g per particle, or 3.4 x 10^8^ g•mol^-1^. This mass is approximately 100 times that of a yeast 80S ribosome (3.2 MDa), consistent with the relative size of stress granule cores (Figure 2C). While the mass of protein-bound mRNAs (mRNPs) is likely to be highly variable between different RNA species and the individual history of the mRNP, it has been estimated^56^ that a “minimal” 1 kb mRNP may contain between 0.3 and 1.3 MDa of protein (plus 0.3 MDa of mRNA), and thus a total mass between 0.6 and 1.6 MDa. For a 3 kb minimal mRNP, the range of masses would be 2 MDa to 4 MDa. Thus, this order-of-magnitude calculation suggests that a given stress granule core particle, which contains many proteins other than those of a minimal mRNP, is likely to contain only a few to a few dozen mRNA molecules. Since our transcriptomic analysis indicates that several thousand different mRNA species are present in the pool of yeast or mammalian stress granule cores, it follows that stress granule cores are likely to be heterogeneous in mRNA composition, paralleling their heterogeneous physical appearance.

To test this hypothesis, we took advantage of the suitability of HEK293T cells for light microscopy. We first imaged the canonical stress granule marker G3BP (after induction of stress granules) using either of two approaches. First, by conventional primary and secondary antibody labeling (Figures 5A and 5B) immunofluorescence (IF), and second, by the hybrid chain reaction (HCR) fluorescence technique (Figures 5A and 5C). The latter overcomes signal diffusion and allows imaging at significantly higher resolution and sensitivity^57^. Consistent with the superior resolution of the more recent technique, stress granules in HEK293T cells appeared either as large diffuse aggregates by the conventional IF technique (Figure 5B) or collections of discrete fluorescence peaks by the HCR method (Figure 5C). These experiments show that HCR-IF can resolve the conventional condensate-like appearance of G3BP in stress granules into finer substructures, possibly stress granule cores.

**Figure 5.**
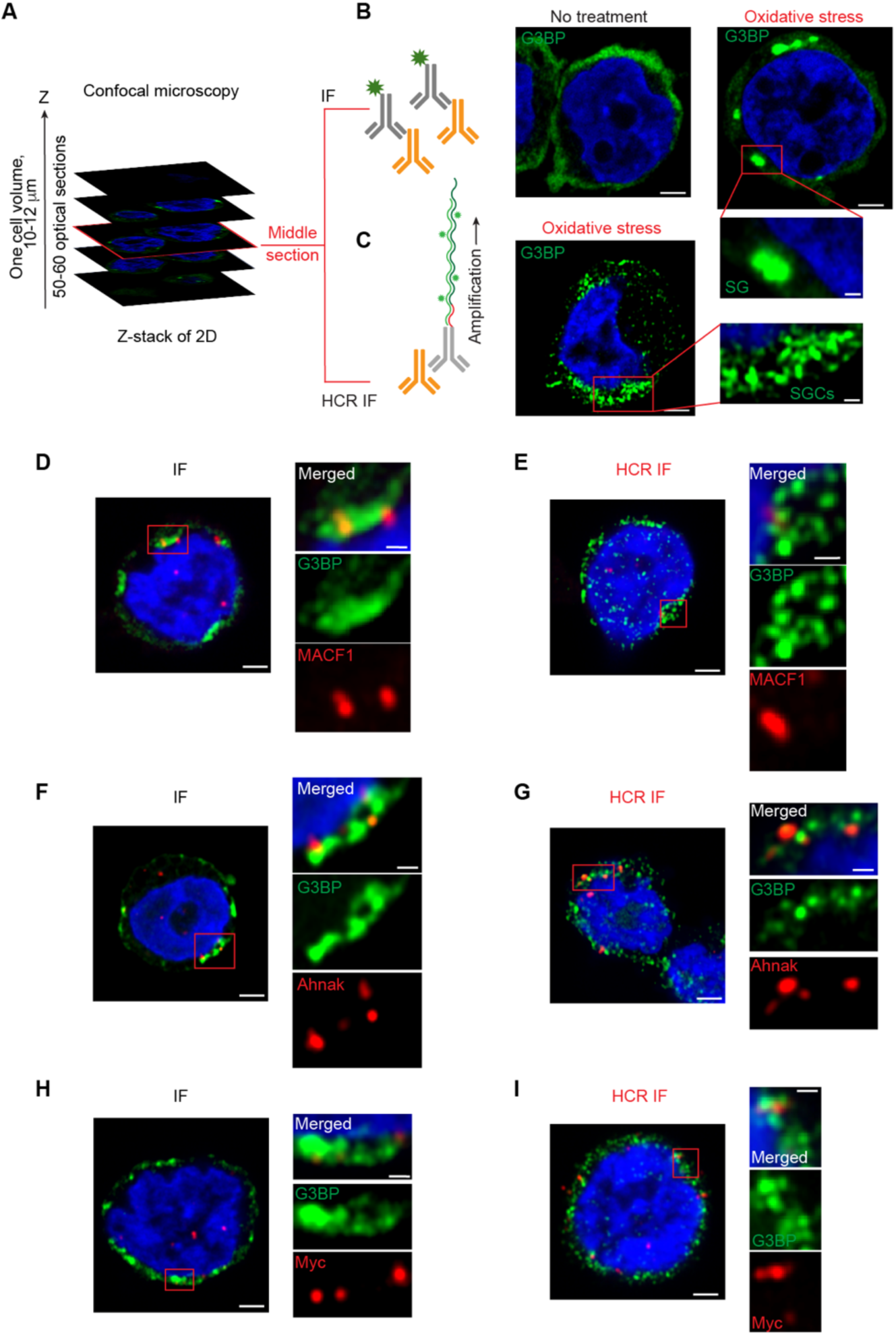
Fluorescent in-situ hybridization analysis of HEK293T cell stress granule cores. (**A**) Schematic of typical Airyscan confocal experiment with Z-stacks covering one cell volume. IF and HCR-IF, immunofluorescence and hybrid chain reaction immunofluorescence, respectively. (**B**) Detection of G3BP (green) by conventional immunofluorescence, with fluorescent label attached to a secondary antibody (grey). Broad G3BP signal corresponding to a stress granule (SG) is framed and magnified. (**C**) Detection of G3BP (green) by the high-resolution HCR approach (schematic). Foci of G3BP signals (SGCs, stress granule cores) are framed and magnified. (**D**, **F**, **H**) Detection of MACF1 (D), AHNAK (F) or MYC (H) mRNA by HCR-FISH and G3BP-labeled stress granules by conventional IF. (**E**, **G**, **I**) Detection of MACF1 (D), AHNAK (F) or MYC (H) mRNA and G3BP by HCR-FISH and HCR-IF. In all panels, unless otherwise indicated, cells underwent oxidative stress. Scale bars 2 μm (full cells) or 0.5 μm (magnified inserts).

We next combined HCR-FISH with either conventional IF (Figures 5D, 5F and 5H) or with HCR IF (Figures 5E, 5G and 5I) to analyze the distribution of specific mRNAs relative to G3BP. First, we examined MACF1 and AHNAK mRNAs, which are enriched in our early log phase HEK293T cell stress granule cores, and have also been reported^22, 58^ to accumulate in stress granules of cells approaching stationary phase. In our experiments using conventional IF, both mRNAs are seen to be embedded in the matrix of stress granules, as defined by the diffuse G3BP signal. In contrast, in our high-resolution HCR-FISH experiments, it is clear that while every signal for both mRNAs colocalizes with a G3BP punctum, many G3BP puncta do not have mRNA signal. Similarly, when identical experiments were carried out for MYC mRNA (Figures 5H and 5I), a global transcriptional promoter whose transcripts are present at a lower titer, a few G3BP puncta colocalized with MYC mRNA as well. The results of these experiments are fully consistent with our hypothesis that stress granule cores are heterogeneous as regards their mRNA composition.

## DISCUSSION

We have developed a new approach to the purification of stress granule cores from yeast and mammalian cells that have been subject to oxidative stress. Our purifications eschew immunoprecipitation, the main enrichment step in previous stress granule purification schemes. While antibodies can provide high levels of enrichment of RNP complexes with simple stoichiometry, their use in purifying large heterogeneous intracellular particles necessarily relies on assumptions about their composition. A case in point is the DNA helicase G3BP1, which is a useful marker for mammalian stress granules (Figure 5), and has been shown to mediate stress granules formation in a dose-dependent manner^59–61^ even in the absence of typical stress^62^. Yet, G3BP1/2 knockout does not lead to complete abrogation of stress granules^63, 64^ and the stress granule transcriptome of such knockout cells remains largely unchanged^58^. Our purification approach, which relies on density and hydrodynamic radius, is unbiased by prior knowledge on the presence or reactivity of particular epitopes, and consequently, can provide information on the compositional heterogeneity of stress granule cores.

An unexpected observation during the course of our purification was the relatively low density of stress granule cores. Our sucrose density-gradient fractionation revealed a stress-induced peak of PABP- or eIF4G1-bound large nucleic acid-containing material with distinctly lower density than ribosomal particles (Figures 1C and 1D, and S1B-S1D). This peak was reduced by cycloheximide treatment and lacked polysomes, consistent with previously reported properties of stress granules, and is composed of particles whose size, stability to high ionic strength, affinity for G-quadruplex binding compounds, and protein and RNA composition are consistent with those of stress granules as determined previously. Our electron microscopy, proteomic and transcriptomic analyses suggest that the relatively low density (for an RNP) of stress granule cores arises from their association with intracellular membranes. This was apparent in negative-stain electron micrographs, especially of the HEK293T cell stress granule cores (Figure 2D and Methods). Our proteomic analyses show that stress granule cores contain components of the endomembrane system including proteins of the endoplasmic reticulum and mitochondria. Previously, it was proposed that the endoplasmic reticulum is the primary site for stress granule biogenesis^65, 66^. Our purification reveals candidate proteins which may provide functional links (Figures 3E and 3G and Tables S2 and S3) such as endoplasmin (HSP90B1, GP96 or GRP94). This is a stress-inducible molecular chaperone, which has a distinct role in innate immunity^50^ and in carcinogenesis^67^. The latter process is tightly linked to stress granule turnover^40, 68^. Contacts between the endoplasmic reticulum and mitochondria are critical to cellular homeostasis^69^. We find that stress granule cores contain proteins that tether these two organelles (*e.g*., GRP75, PARK7, BAP31 and VAPB; Table S3), which may suggest that under stress, the endoplasmic reticulum funnels signals between the stalled translational machinery and the powerhouse of the cell. Hitherto, the association of stress granules with intracellular membranes was mainly deduced from microscopy ^48, 65, 70–72^. Interestingly, intrinsically disordered protein regions play critical roles in many functions of cellular membranes as well as in formation of stress granules^73–75^. Our purifications provide now molecular entry points for studying interconnections between these two categories of cellular structures and delineate a complex global cellular network of interactions involving various pathways (Figure 6 and Tables S8 and S9).

**Figure 6.**
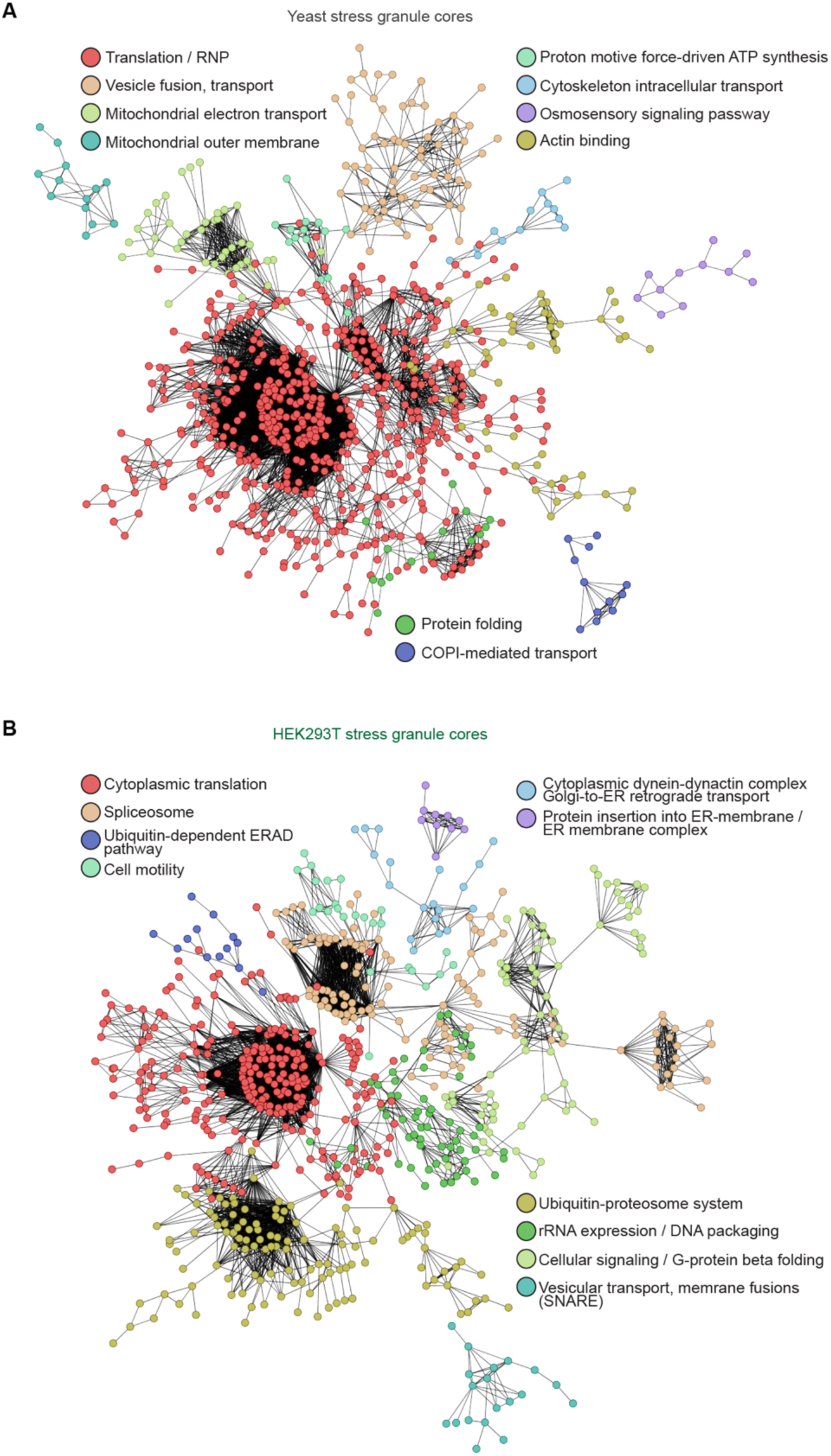
Protein-protein interactomes for yeast and mammalian stress granule cores. (**A**, **B**) A network of interactions within yeast (A) and mammalian (B) stress granule cores differentiated into ten k-means clusters (STRING database^78^, see Methods). Active interaction sources set to ‘Experiments’ mode only. The edges indicate both functional and physical protein associations.

Our purification of stress granule cores from different yeast strains and growth conditions shows that the composition of these RNPs can vary depending on these variables (Tables 1 and S2). Our physical characterization of stress granule cores provides direct evidence of their heterogeneity (Figures 2A and 2B). Moreover, our determination of the density and mean size of stress granule cores yields an approximate mass range for these RNPs, and sets physical constraints on their composition. In particular we find that each particle is too small to contain the entire stress granule core transcriptome, implying that different stress granule cores contain different subsets of mRNAs. Our high-resolution FISH analysis of stress granules in HEK293T cells supports this, showing that while the examined mRNAs enter stress granules, not all stress granules are associated with particular mRNAs. Recent evidence suggests that stress granules assemble from nanoscale seeds that exist in the absence of stress^4, 54, 64, 76, 77^. Such sub-microscopic seeds would be heterogeneous (as they would be too small to contain all mRNAs), and their composition may determine that of the stress granule cores they nucleate. Our physical purification of stress granule cores now opens the way to the biochemical analysis of their compositional and functional diversity and their assembly pathways.

## Supporting information

Main Table 1

## ACKNOWLEGEMENTS

We are grateful to Drs. G. Piszceck and D. Wu of the Biophysics Core of the National Heart, Lung, and Blood Institute (NHLBI) for assistance in characterization of the purified stress granule cores; Y. He from the fermentation facility (NHLBI) for growing HEK293T cell cultures; Dr. X. Wu of the Light Microscopy Core (NHLBI) for confocal microscopy support; the staff of the Electron Microscopy Core (NHLBI) for technical assistance; Dr. M. Gucek and S. Patel of the Proteomics Core (NHLBI) for mass spectrometric analyses; Dr. C. Jones and all other members of the Ferré-D’Amaré laboratory for discussions. This work was supported by the intramural program of the NHLBI, National Institutes of Health, USA.

## AUTHOR CONTRIBUTIONS

Study conceived and designed by N.A.D. and A.R.F.-D. Experiments carried out by N.A.D. Data analyzed by N.A.D. and A.R.F.-D. Manuscript written by N.A.D. and A.R.F.-D.

## METHODS DETAILS

### Yeast strains and plasmids

Yeast strains with chromosomally encoded GFP fusions were purchased from the Yeast GFP Clone Collection (Thermo Fisher Scientific). Strains JD1370 (*MATalpha trp1 ura3 leu2 [L-A] [Mi] PEP4::HIS3 NUC1::LEU2*), and YAG1021 (*MATalpha W303 hht2::FLAG-HHT2-URA3*), were kindly provided by Professors J. Dinman (University of Maryland, College Park, MD, USA) and A. Gunjan (Florida State University, Tallahassee, FL, USA), respectively. HEK293T cells were purchased from Millipore Sigma (96121229-1VL).

### SDGC-SEC purification of *S. cerevisiae* stress granule cores

A typical isolation started with 6 L of culture. Yeast grown in YPD with adenine (YPDA) or synthetic defined (SD) media supplemented with 0.05 g/L kanamycin to an OD_600_ of 0.5-0.65 were treated with 0.5% (w/v) NaN_3_ for 40-45 min at 30 °C while shaking at 225 rpm. Cells were collected by centrifugation at 4000 × *g* for 3 min at 30 °C in prewarmed bottles, then resuspended in 50 mL of the same media, centrifuged at 4000 × *g* for 3 min at 30 °C, and immediately frozen in liquid nitrogen. Stressed YER165W Pab1-GFP or JD1370 cells yielded 0.8-0.9 g wet weight per L of culture. All following procedures were performed at 3 °C. Cells were thawed on ice during 5 min, resuspended in buffer 5/125, centrifuged at 4000 × *g* for 3 min at 2 °C, then resuspended in the same buffer again (1 mL buffer/g of cells) supplemented with sodium heparin (Sigma), AEBSF (EMD Millipore Corp.), RNasin Plus (Promega), SUPERase-In (Invitrogen), and EDTA-free protease inhibitor cocktail (PIC; Sigma-Aldrich). Lysis was performed manually by vortexing with prechilled (-20 °C) glass beads (3 g of acid-washed 425-600 μm beads/g of cells) for six one-minute cycles interspersed with one-minute incubations on ice. Lysed cells were centrifuged at 850 × *g* for 2 min, then supernatant was centrifuged at 900 × *g* for 2 min and further at 18000 × *g* for 20 min. The resultant viscous supernatant (1 g of cells yielded ∼ 0.8 mL with A_260_ of 160 – 200) was supplemented with inhibitor of proteases and RNases again, and immediately loaded on a chilled (3 °C) sucrose density gradient prepared in buffer 4/125, and centrifuged at 29,000 rpm for 13.25 h (SW 32 Ti, Beckman). Aliquots of sucrose density gradient centrifugation (SDGC) fractions were mixed with loading buffer (1 × TAE, 0.1% (w/v) BPB, 30% glycerol (v/v), 34% (w/v) sucrose) and assayed by agarose gel electrophoresis [1% (w/v), 1 × TAE, 1 μg/mL ethidium bromide]. Fractions with large nucleic acids were pooled and centrifugally concentrated at 3200 × *g* at 3 °C. The sample was diluted with buffer 2.5/100 and then urea was added to final concentrations of 2 M. After incubation at 3 °C, the sample was loaded on a HiPrep 16/60 Sephacryl S-500 column (Cytiva) equilibrated in buffer 2.5/100 with or without 2M urea. Size-exclusion chromatography (SEC) was carried out at 0.25 mL/min. Fractions enriched in stress granule cores (assayed by 1% agarose gel electrophoresis as above) were centrifugally concentrated to 8-10 A_260_ units/mL at 3 °C and used immediately or flash-frozen in liquid nitrogen and stored in small aliquots at -80 °C (thawed material was never refrozen).

### SDGC-SEC purification of mammalian stress granule cores from HEK293T cells

Each isolation used HEK293T derived tsA201 cells eradicated from mycoplasma at ECACC. The identity of tsA201 and 293 has been confirmed by STR profiling. A typical isolation started with 8 L of HEK293T cells. One frozen vial with 1 × 10^7^ tsA201 cells was inoculated to 40 mL of Gibco FreeStyle 293 Expression Medium with 2% Fetal Bovine Serum, 50 U/mL Penicillin-Streptomycin and 2 mM L-Glutamine in a 250 mL flat bottom cell culture shaker flask. Cells were grown for three days in Infors Multitron incubator at 130 rpm and 75 % humidity with 8 % (v/v) CO_2_. Cells were then passed into a large flask (2 or 3 L Fernbach Flask, Vent Cap, Plain Bottom) with addition of above mentioned medium to reach a final cell density of 0.8 -1.0 × 10^6^. Cells were treated with 0.5 mM sodium arsenite (LabChem) for 1 h and immediately collected in bottles pre-warmed to 37 °C at 300 × *g* for 10 min. Cells were then washed in ice-cold buffer 5/150, centrifuged at 300 × *g* for 9 min at 4 °C, resuspended in the same buffer 5/150 with 2 mM DTT (0.7 mL buffer/g of cells; see Figure 1B) supplemented with heparin, AEBSF, RNasin Plus, SUPERase-In and PIC as for yeast. Plasma membranes were immediately disrupted by decompression on ice at 1000 psi (Parr Instrument Company), then the sample was centrifuged at 850 × *g* for 2 min at 4 °C with following spinning of supernatant at 900 × *g* for 2 min at 4 °C. The resultant supernatant was centrifuged at 18,000 × *g* for 15 min at 4 °C, then NP-40 was added to 0.5% w/v and the lysate was incubated on ice. The material was then loaded on chilled (3 °C) sucrose density gradients and centrifuged overnight in rotor SW 32 Ti (see yeast protocol). The sucrose gradients were fractionated as described above. Top fractions were combined (see Figure S2B) and centrifugally concentrated at 3200 × *g* at 3 °C. The sample was diluted and then urea was adjusted to 2 M. After incubation at 3 °C, the sample was loaded on a 16/60 Sephacryl S-1000 column equilibrated in a buffer with 2 M urea. SEC was carried out at 0.35 mL/min (Figure S2C). Fractions enriched in large molecular weight nucleic acids were centrifugally concentrated as above to 2 A_260_ units/mL at 3 °C and used immediately or flash-frozen in liquid nitrogen and stored in small aliquots at -80 °C (thawed material was never refrozen).

### Purification of yeast 80S ribosomes

Yeast cells (strain JD1370) were grown and stressed as above in YPD with adenine and kanamycin and used for purification of the 80S ribosomes as described^88^.

### Nanoparticle Tracking Analysis (NTA)

SDGC-SEC-purified stress granule cores from yeast were diluted in buffer 2.5/100 with 0.5 mM DTT (urea was adjusted to 80 mM) and analyzed using the ZetaVIEW Nano Particle Analysis system (Analytik). The laser was set at 660 nm (n=3). A representative acquired track was 7121 traces with 592 average counted particles per frame (Figure 2A). SDGC-SEC-purified stress granule cores from HEK293T cells were analyzed similarly using buffer 2.5/80/40 with 0.5 mM DTT (urea was adjusted to 100 mM) and laser of 488 nm to detect particles that fall in the smaller size range. On average 1000 tracks were acquired with 200 particles per frame (n=5). Data were analyzed using ZetaView 8.05.04 software (www.particle-metrix.de).

### Negative-stain electron microscopy

SDGC-SEC-purified stress granule cores from yeast (strain YAG1021) were diluted in buffer 2.5/100 without EDTA to 2.0 A_260_ units/mL, incubated at 37°C for 2h, and applied on glow-discharged grids (Electron Microscopy Sciences, FCF300-CU Formvar/Carbon Mesh, Copper; 50 sec at 21 °C). To prevent stress granule cores fusion the grids were blotted and treated in 0.1% CWFS gelatin (Aurion) in Tris-buffered saline (TBS) at pH 7.6 for 30 min at 4 °C. Then, the grids were re-blotted and incubated with primary antibodies (Millipore-Sigma, Cat#F1804) used at a dilution of 1:30 in TBS for 50 min at 21 °C. Grids were washed three times in TBS for 5 min at 21 °C, and treated with secondary antibodies (Nanoprobes, Cat#2001) used at a dilution of 1:50 to 1:100 in TBS with 0.1% CWFG for 45 min at 4 °C. Grids were washed three times as above and stained with freshly prepared 1% (w/v) uranyl formate for 90 sec, and then re-stained for another 50 sec. Images were acquired on a FEI Tecnai T12 transmission electron microscope (Thermo Fisher Scientific). In parallel, grids with yeast 80S ribosomes at A_260_ of 0.25 units/mL were stained with 1% (w/v) uranyl formate for 40 sec at 21 °C and electron micrographs were obtained. Negative-stain experiments for the HEK293T stress granule cores were carried out by pretreatment of the SDGC-SEC-purified samples with n-dodecyl-b-maltoside taken at one or two critical micelle concentration in “buffer 2.5/80/40” for 1 h at 4 °C. The grids were stained with uranyl formate for 90 sec; the images were acquired as above.

### Enrichment of yeast stress granules by immunoprecipitation

6 L of yeast culture YER165W-GFP-Pab1 grown in YPD with adenine and kanamycin was stressed and then split equally. For comparison of proteomes (Figure 3A), yeast stress granule cores were either purified as described^17, 21^ with proportional scaling of all steps to 3 L culture, or by the SDGC-SEC protocol above.

### Proteomics

For analysis of yeast stress granule cores (Tables 1 and S2; datasets 1-4), 1.0 A_280_ unit of the cores purified from strains YER165W Pab1-GFP or JD1370 was incubated in 1.5 – 2% (w/v) SDS for 30 min at 55 °C with shaking at 300 rpm. Total proteins were extracted with acetic acid, and then precipitated with acetone, as described^89, 90^. Pellets of 0.1 mg total protein were further subjected to standard in-solution digestion (disulfide reduction, alkylation and trypsin digestion). To minimize missed cleavage sites, digestion with rLys-C was carried out prior to trypsin treatment. Proteins extracted from the immunoprecipitation-enriched stress granule preparation^17^ were treated identically. Tryptic peptide analysis was performed using an Orbitrap Fusion Lumos Tribid mass spectrometer (Thermo Fisher Scientific) interfaced with an Ultimate 3000 Nano-HPLC (Thermo Fisher Scientific). Peptides were fractionated on an EASY-Spray PepMAP RPLC C18 column (2 μm, 100Å, 75 μm x 50 cm) with a 120 min, 5 – 35% (v/v) linear acetonitrile gradient [with 0.1% (v/v) formic acid] at a flow rate of 300 nL/min. The instrument was operated in data-dependent acquisition mode (DDA) using the FT mass analyzer for one survey MS scan on selecting precursor ions, followed by 3 second data-dependent HCD-MS/MS scans for precursor peptides with 2-7 charged ions above a threshold ion count of 10,000 with normalized collision energy of 37%. Survey scans of peptide precursors from 300 to 2000 m/z were performed at 120k resolution and MS/MS scans were acquired at 50,000 resolutions with a range of m/z 100-2000. All MS and MS/MS raw spectra from each set were processed and searched using the Sequest HT algorithm within Proteome Discoverer 2.2 software (PD2.2 Thermo Fisher Scientific). The settings for precursor mass tolerance were set at 12 ppm, fragment ion mass tolerance to 0.05 Da, trypsin enzyme with 2 missed cleavages with carbamidomethylation of cysteine as fixed modifications, deamidation of glutamine and asparagine, oxidation of methionine as variable modifications. The SWISS-PROT *S. cerevisiae* sequence database was used for the search. Identified peptides were filtered for maximum 1% FDR using the Percolator algorithm in PD 2.2 along with additional peptide confidence set to medium. The final lists of protein identification/quantitation were filtered by PD 2.2 (identification set at medium confidence). For the quantitation, a label-free approach was used, where the area under the curve for the precursor ions was used to calculate the relative fold change between different peptide ions.

Analysis of the proteins from yeast and mammalian stress granule cores was also carried out by Poochon Scientific (yeast: Tables 1 and S2; datasets 5 and 6; mammalian: Tables 2 and S3). Samples were processed for trypsin/LysC digestion as per Standard Operation of Procedure for in Solution Digestion (SOP-PS-6003). The digested peptide mixture was then concentrated and desalted using C18 Zip-tip as per Standard Operation of Procedure for Desalting Digested Peptides (SOP-PS-6005). Desalted peptides were reconstituted in 0.1% formic acid and each sample was analyzed by 110 min LC/MS/MS run. The LC/MS/MS analysis of tryptic peptides for each sample was performed sequentially with a blank run between each two sample runs using a Thermo Scientific Orbitrap Exploris 240 Mass Spectrometer and a Thermo Dionex UltiMate 3000 RSLCnano System. Peptides from trypsin digestion were loaded onto a peptide trap cartridge at a flow rate of 5 μL/min. The trapped peptides were eluted onto a reversed-phase Easy-Spray Column PepMap RSLC, C18, 2 μM, 100A, 75 μm × 250 mm (Thermo Scientific) using a linear gradient of acetonitrile (3-36%) in 0.1% formic acid. The elution duration was 110 min at a flow rate of 0.3 μl/min. Eluted peptides from the Easy-Spray column were ionized and sprayed into the mass spectrometer, using a Nano Easy-Spray Ion Source (Thermo Scientific) under the following settings: spray voltage, 1.6 kV, Capillary temperature, 275°C. Other settings were empirically determined. Raw data files were searched against human protein sequences database using the Proteome Discoverer 3.0 software (Thermo, San Jose, CA) based on the SEQUEST algorithm. Carbamidomethylation (+57.021 Da) of cysteines was set as fixed modification, and Oxidation / +15.995 Da (M), Phospho / +79.966 Da (S, T, Y), Acetylation/+41.011 Da (K), and Deamidation / +0.984 Da (N, Q) were set as dynamic modifications. The minimum peptide length was specified to be five amino acids. The precursor mass tolerance was set to 15 ppm, whereas fragment mass tolerance was set to 0.05 Da. The maximum false peptide discovery rate was specified as 0.01. To compare the proteins in abundance among samples the data were normalized using the normalized spectral abundance factors method to calculate the protein relative abundance^91, 92^. Venn diagrams were created using http://bioinformatics.psb.ugent.be/webtools/Venn/. The GO enrichment analysis was performed using https://geneontology.org. Protein-protein interactions for top 2000 genes filtered by the highest coverage were analyzed by STRING database^78^ by applying ‘Experiments’ evidence mode with interconnected nodes defined by highest confidence (set at 0.85 for yeast and at 0.9 for HEK293T) and one interaction per first shell. Figure 6 represents such STRING analysis for yeast dataset 5 (Table S2) and HEK293T dataset 3 (Table S3). First, 2000 genes were analyzed by clustering using 99 and 72 k-means for yeast and HEK293T datasets, respectively. This analysis allowed the identification of one dominant cluster per each dataset, which was further re-clustered into 10 k-means (Figure 6 and Tables S8 and S9).

### RNA Seq

Yeast samples were firstly treated with RNase-free proteinase K (Invitrogen) for 1 h 30 min at 50 °C with moderate shaking. Then, RNA was extracted by using TRIzol reagent (Ambion) according to manufacturer’s protocol with additional heating at 60 °C for 20 min. The top 40% of the aqueous phase was used for downstream analysis. RNA was precipitated using RNase-free glycogen (0.25 g/L final) as a carrier by addition of equal volume of isopropanol. Prior to sequencing RNA was treated with DNase (Invitrogen). Sequencing and analysis were performed by Novogene and Genewiz/Azenta using the Illumina platform with paired end reads. Prior to library creation ribosomal RNA was removed using Illumina Ribo-Zero Depletion or QIAseq FastSelect -rRNA Yeast kits.

For RNA from stress granule cores of *S. cerevisiae* grown in YPD with adenine and kanamycin, qualified trimmed reads were aligned to the reference genome (iGenomes, Ensembl R64-1-1) using HISAT. Distribution of reads within known RNA types was carried out by HTseq. Assembly was performed by the StringTie package (v1.3.1). Identification of lncRNA was done by screening length and known annotations, and by coding potential using Pfam-scan (v1.3; -E 0.001 --domE 0.001 -pfamB) and CPC (cpc-0.9-r2; e-value = 1e-10). Quantification of mRNAs, lncRNAs and TUCPs was assessed by StringTie-eB. For RNA from crude lysates (‘SUPERNATANT’, 18000 × *g*, see Figure 1A) and from stress granule cores of *S. cerevisiae* grown in SD medium, qualified trimmed reads were mapped to the reference genome (iGenomes, sacSer3) using the STAR aligner (v.2.5.2b). Unique gene hit counts were calculated by using feature Counts from the Subread package (v.1.5.2). Only unique reads that fell within exon regions were counted. Differentiation analysis was performed by using DESeq2. The Wald test was used to generate p-values and Log2 fold changes. Genes with adjusted p-values < 0.05 and absolute log2 fold changes > 1 were called as enriched and depleted for each comparison. Samples of RNA from crude lysates (‘SUPERNATANT’, 18000 × *g*, see Figure 1B) and from stress granule cores of HEK293T cells were obtained as described above. Prior to sequencing RNA was treated with DNase (Invitrogen) and ribosomal RNA was removed using Illumina Ribo-Zero Plus kit. Sequencing and analysis were performed by SeqCenter using the Illumina platform with paired end reads. Read mapping was performed via STAR to the hg38 version of the human genome (GRCh38.p14). Feature quantification was performed using RSEM. Normalization was carried out by using edgeR. Subsequent values were then converted to counts per million. Differential analysis was performed using edgeR’s exact test (exactTest) for differences between two groups of negative-binomial counts with an estimated dispersion value of 0.1. Differentially varying genes (enriched versus depleted) were determined as above (adjusted p-values < 0.05; log2 fold changes > 1). Gene ontology for yeast and HEK293T datasets was performed on the statistically significant set of genes by using the Gene Ontology Resource (https://geneontology.org).

### Analysis of yeast stress granule cores stability in high ionic strength

SDGC-SEC-purified stress granule cores (strain JD1370) or yeast 80S ribosomes were incubated at 0.6 and 3.0 A_260_ units/mL, respectively, in buffer 2/150 supplemented with either 1.0 M KCl, 1.0 M CsCl, 50 mM EDTA or 1.0 M LiCl for 1 h at 30 °C. Controls were incubated in unsupplemented buffer 2/150. Samples were analyzed as described above except using a longer electrophoretic run to resolve 60S and 40S ribosomal subunits.

### Interaction with G-quadruplex-binding compounds

SDGC-SEC-purified stress granule cores from yeast (strain JD1370) were incubated at 0.3 A_260_ units/mL in buffer 2.5/120 in the presence of 0.05, 0.1, 0.5, 1, 2 and 10 μM pyridostatin (PDS; Tocris Bioscience) for 30 min at 30°C. To investigate the combinatorial effect of PDS and LiCl, the stress granule cores were first incubated with 1.0, 5.0 or 10.0 μM PDS for 20 min at 30°C; then LiCl was added to 1M, and the treatment continued for an additional 30 min. To investigate the combinatorial effect of PDS and N-methyl mesoporphyrin IX (NMM; Frontier Scientific), stress granule cores were incubated with PDS at 5.0 μM and NMM at 0, 1.0, 2.0, 5.0, 10.0 or 20 μM in buffer 2.5/120 for 30 min at 30 °C. All samples were analyzed by electrophoresis in 1% agarose gels as above.

### Fluorescent microscopy of yeast spheroplasts

Yeast strains with endogenous fusions of GFP (eIF4A, ABF1 and RAP1) were cultured in SD medium with 1.5% w/v D-glucose and 0.75% w/v D-(+)-galactose to an OD_600_ of 0.4-0.5 and treated with 0.5% w/v NaN_3_ for 40 min at 30 °C while shaking at 225 rpm. Cells were then fixed with 4.5% (v/v) formaldehyde for 40 min at 21 °C while shaking at 100 rpm, and collected by centrifugation at 4500 × *g* for 10 min at 21 °C. Yeast pellets were washed with ice-cold 100 mM K-phosphate, pH 6.5, incubated on ice for 10 min and re-centrifuged at 4500 × *g* at 4 °C. The washing procedure was repeated once. Cells were then resuspended in ice-cold 1.2M Sorbitol in 100 mM potassium phosphate, pH 6.5 and supplemented with zymolyase and 2-mecraptoethanol as per manufacture instructions (ZymoResearch E1005). Cell walls were digested for 30 min at 30 °C while shaking at 300 rpm. Spheroplasts were collected by centrifugation at 400 × *g* for 15 min at 21 °C, then washed in sorbitol buffer, span as above, dissolved in the sorbitol buffer and applied on poly-L-Lysine (1 mg/mL) covered slides for overnight at 4 °C. Permeabilization was carried out in 70% ethanol at -20 °C for 2 h 25 min. Immobilized spheroplasts were rehydrated in wash buffer (WB; 2 × SSC, 10% v/v deionized formamide) for 5 min at 21 °C and incubated with 125 nM T30-ATTO 647N (Biosearch Technologies) in hybridization buffer (WB with 10% w/v dextran sulfate and 0.02 mg/mL BSA) for 4 h 15 min at 37 °C in the dark. Slides were washed with WB prewarmed to 37 °C for 30 min in the dark, the wash was repeated once. Finally, slides were rinsed with 2 × SSC at 21 °C and then incubated with 1 μg/mL DAPI in 1 × PBS for 5 min at 21 °C. Fluorescence images were acquired with a Zeiss LSM880 Airyscan confocal microscope (SR-mode) using Plan-Apochromat 63×/1.4 Oil DIC M27 objective. About 25 Z-stacks (0.15-0.2 μm depth; one spheroplast volume is ∼ 4-5 μm) were taken from each visual field. Airyscan processing was done by ZEN Black software in 3D mode (automatic setting). Images in Figures 3B-3D are one middle Z-stack; display values were optimized

### Hybridization chain reaction-fluorescence in situ hybridization and Immunofluorescence

Aliquots of HEK293T cells (5-30 mL) grown in suspension for SDGC-SEC purification (see above) were taken directly before addition of 0.5 mM sodium arsenite and fixed in 3.7% formaldehyde in 1 × PBS, pH 7.4 for 10 min at 21 °C. Cells treated with oxidative stress for 1 h at 37 °C were fixed identically. Untreated and stressed fixed cells were washed 3 times in 1 × PBS and used immediately or stored at -20 °C in 70% (v/v) ethanol. Fixed cells were immobilized on poly-L-Lysine covered eight-well chamber cover glass (#1.5 high performance, Cellvis). IF and HCR-IF were carried out as specified below by using HCR-FISH probes. HCR-FISH probes for detection of MACF1 (chr1: 39,387,374-39,387,965), AHNAK (chr11: 62,532,826-62,533,957) and MYC (chr8: 127,736,138-127,736,621 and 127,740,450-127,741,041) mRNA targets, and secondary antibody for HCR-IF were ordered from Molecular Instruments, Inc. The hybridization step with FISH probes was followed as per vendor’s protocol. The following immunostaining step was performed using mouse anti-G3BP (1:300; Abcam ab56574) for 1 h at 21 °C followed by three washes with 1 × PBS, pH 7.4 for 5 min each. Slides were then incubated with either goat anti-mouse FITC antibody (1:700; Abcam ab6785; Figures 5B, 5D, 5F and 5H) or donkey anti-mouse secondary antibody with an initiator for use with amplifier B1-488 (1:700; Molecular Instruments; Figures 5C, 5E, 5G and 5I) for 1 h at 21 °C and washed three times with 1 × PBS, pH 7.4 for 5 min each. The amplification step was followed as per vendor’s protocol using amplifiers B2-546 for detection of RNA and B1-488 for staining G3BP. Samples were covered with ProLong Glass antifade mounting media with 0.6 μg/mL Hoechst 33342. Fluorescence images were acquired with a Zeiss LSM880 Airyscan confocal microscope as above. Approximately 50-60 Z-stacks (0.2 μm depth) were taken from each visual field. Airyscan processing was done by ZEN Black software in 3D mode (automatic setting). Images in Figure 5 are one middle Z-stack; display values were optimized.

## Materials availability

Requests should be submitted by email to A.R.F.-D. Per manufacturer’s policies, HCR-FISH probes can be requested from Molecular Instruments, Inc. referring to lots RTK099, RTG842 and RTG844 for MACF1, AHNAK and MYC mRNA targets, respectively.

## Data availability

The mass spectrometry proteomics data have been deposited to the ProteomeXchange Consortium via the PRIDE^93^ partner repository with the dataset identifier PXD053088 (*S. cerevisiae* strain JD1370 grown in rich medium), PXD053090 (*S. cerevisiae* strain JD1370 grown in SD medium), PXD053260 (*S. cerevisiae* strain YER165W with PABP1-GFP fusion grown in rich medium) and PXD053091 (HEK293T grown in suspension). Raw sequencing data have been deposited in the Sequence Read Archive. Data for yeast are deposited under BioProject PRJNA941899 (YPDA medium, stress granule cores: SRR23719788 and SRR23719789; SD medium, ‘18000 × *g*’ lysates: SRR29249735 and SRR29249734; SD medium, stress granule cores: SRR29249737 and SRR29249736). Data obtained from HEK293T cells are deposited under BioProject: PRJNA1018544 (‘18000 × *g*’ lysates: SRR29266359, SRR29266360 and SRR29266361; stress granule cores: SRR29264088, SRR29264089 and SRR29264090).

