## Supplemental Figures S1-S3 for "New purifications reveal yeast and human stress granule cores are discrete particles with complex transcriptomes and proteomes"

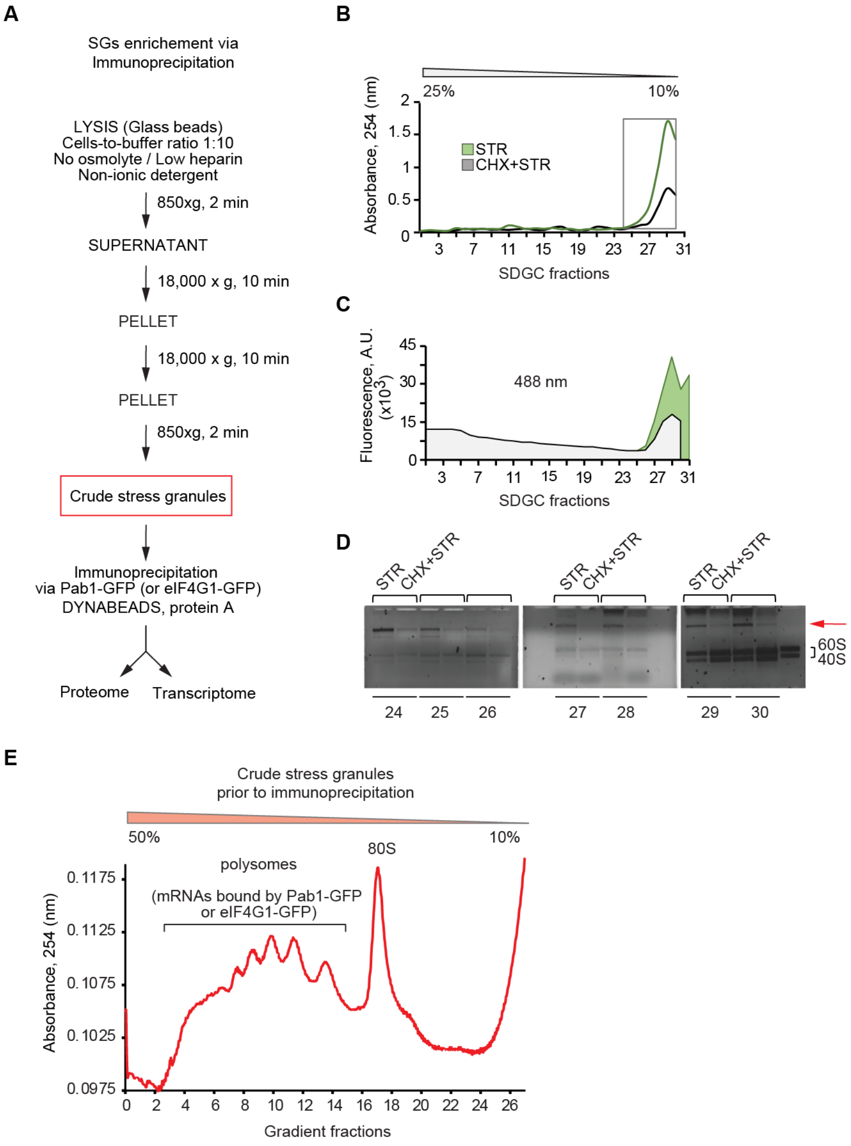


**Figure S1. Immunoprecipitation protocol for purification of stress granules from *S. cerevisiae*.** (**A**) Schematic of stress granule enrichment by the immunoprecipitation-based method[1,2]. (**B**) Analysis of the crude stress granules fraction (red frame in (A)) by narrow range 10-25% (w/v) sucrose density gradient centrifugation (SDGC). 'STR' and 'CHX+STR', material derived from stressed cells and from cells treated with cycloheximide (CHX) prior to stress. Strain used is with endogenous eIF4G1-GFP fusion. (**C**) Analysis of GFP fluorescence of the top SDGC fractions from (B). (**D**) Electrophoretic analysis of the SDGC fractions from (B) (red arrow, large nucleic acids in the eIF4G1-GFP-enriched fractions). Electrophoretic mobility of ribosomal subunits noted. The control ribosome (right) lane was loaded with 80S ribosomes that dissociate into subunits during electrophoresis, as reflected by slightly slower mobility. (**E**) Analysis of the crude stress granules fraction obtained through the immunoprecipitation protocol[1,2] (red frame in (A)) by wide range 10-50% (w/v) SDGC.

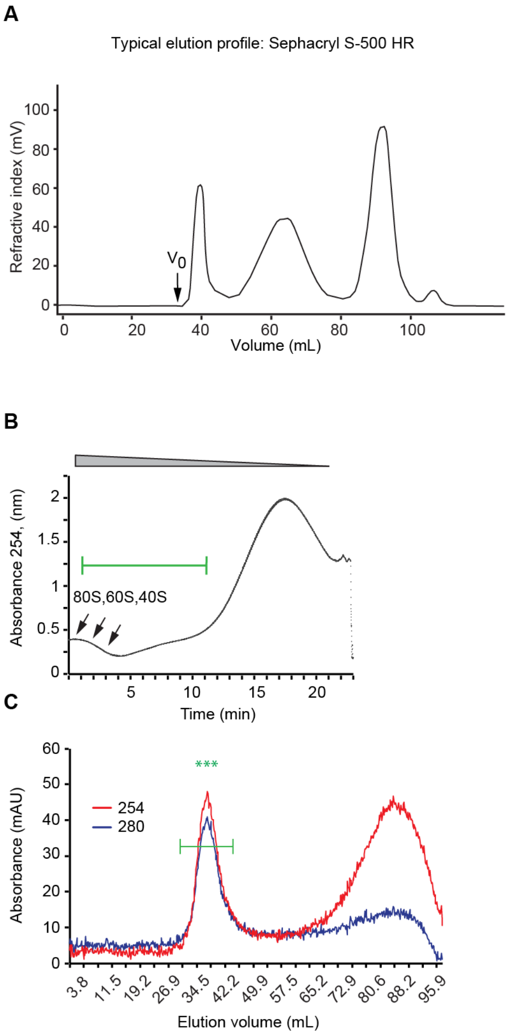


**Figure S2. SDGC-SEC protocol for purification of stress granule cores from yeast and mammalian cells.** (**A**) Elution profile for size-exclusion liquid chromatography column (Sephacryl S-500 HR 16/60; Cytiva) used for stress granule cores purification from yeast. $V_0$, void volume. (**B**) Exemplary SDGC profile of cytoplasmic fraction of HEK293T cells exposed to oxidative stress. Green line, analyzed fractions. (**C**) Exemplary elution profile for size-exclusion liquid chromatography column (Sephacryl S-1000 SF 16/60; Methods) used for stress granule cores purification from HEK293T cells.

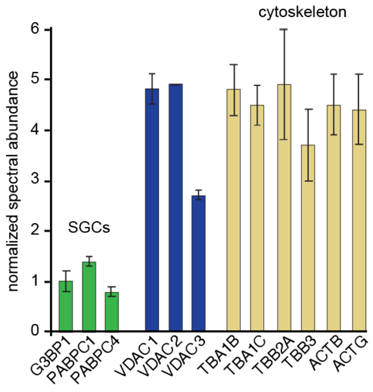


**Figure S3. Proteome characteristics for stress granule cores purified by SDGC-SEC protocol from HEK293T cells.** Illustrative abundant stress granule cores proteins from HEK293T cells (n=2; average ± SD). Relative abundance of proteins between datasets was normalized using G3BP1 (see Methods).

**Supplemental Tables**

| | |
|---|---|
| Table S1. | Overlap of proteomes for yeast stress granules enriched by immunoprecipitation and for stress granule cores isolated by SDGC-SEC protocol (this study). |
| Table S2. | Proteomes for yeast stress granule cores isolated by SDGC-SEC protocol. |
| Table S3. | Proteomes for HEK293T stress granule cores isolated by SDGC-SEC protocol. |
| Table S4. | Transcriptomes for yeast stress granule cores isolated by SDGC-SEC protocol. Part 1. |
| Table S5. | Transcriptomes for yeast stress granule cores isolated by SDGC-SEC protocol. Part 2. |
| Table S6. | Transcriptomes for HEK293T stress granule cores isolated by SDGC-SEC protocol. Part 1. |
| Table S7. | Transcriptomes for HEK293T stress granule cores isolated by SDGC-SEC protocol. Part 2. |
| Table S8. | k-means clustering of protein-protein interactome for yeast stress granule cores isolated by SDGC-SEC protocol. |
| Table S9. | k-means clustering of protein-protein interactome for mammalian (HEK293T cells) stress granule cores isolated by SDGC-SEC protocol. |
